# cdsBERT - Extending Protein Language Models with Codon Awareness

**DOI:** 10.1101/2023.09.15.558027

**Authors:** Logan Hallee, Nikolaos Rafailidis, Jason P. Gleghorn

## Abstract

Recent advancements in Protein Language Models (pLMs) have enabled high-throughput analysis of proteins through primary sequence alone. At the same time, newfound evidence illustrates that codon usage bias is remarkably predictive and can even change the final structure of a protein. Here, we explore these findings by extending the traditional vocabulary of pLMs from amino acids to codons to encapsulate more information inside CoDing Sequences (CDS). We build upon traditional transfer learning techniques with a novel pipeline of token embedding matrix seeding, masked language modeling, and student-teacher knowledge distillation, called MELD. This transformed the pretrained ProtBERT into cdsBERT; a pLM with a codon vocabulary trained on a massive corpus of CDS. Interestingly, cdsBERT variants produced a highly biochemically relevant latent space, outperforming their amino acid-based counterparts on enzyme commission number prediction. Further analysis revealed that synonymous codon token embeddings moved distinctly in the embedding space, showcasing unique additions of information across broad phylogeny inside these traditionally “silent” mutations. This embedding movement correlated significantly with average usage bias across phylogeny. Future fine-tuned organism-specific codon pLMs may potentially have a more significant increase in codon usage fidelity. This work enables an exciting potential in using the codon vocabulary to improve current state-of-the-art structure and function prediction that necessitates the creation of a codon pLM foundation model alongside the addition of high-quality CDS to large-scale protein databases.

## 1 Introduction

Understanding how a protein’s sequence impacts its overall function, physiochemical properties, efficacy, and stability is vital for deciphering mechanisms that underpin biology, including replication, transcription, translation, metabolism, molecular signaling, and even disease-state-specific interaction networks [1, 2, 3, 4, 5, 6]. Protein Language Models (pLMs) have taken biomedical research by storm, allowing for unprecedented large-scale protein analysis. Some notable contributions include AlphaFold2, RoseTTAFold, ESMFold, OmegaFold, and EMBER2, which have successfully estimated amino acid sequence-to-structure mapping [7, 8, 9, 10, 11]. More generalized models such as ProtBERT, ProtT5, Ankh, and xTrimoPGLM offer highly effective contextualized sequence representations that map intuitively to protein function, gene ontology, physiochemical properties, and more [12, 13, 14]. Interestingly, some pLM projects have opted for different vocabularies outside of the traditional single-letter amino acid code. For example, the generative model ProtGPT2 employs a tokenizer encompassing over 50,000 recurrent common oligomers, enhancing the scope of the token embedding matrix (TEM) by introducing more vocabulary variety [15]. However, the existence of codons within biology presents a promising biologically pertinent vocabulary to use for token embedding.

At the core of molecular biology lies the central dogma, which encapsulates the flow of genetic information from DNA to RNA to proteins [16]. Codons, comprising three nucleotides for every one amino acid, serve an intermediary role in this process, encoding the 20 standard amino acids as well as three distinct stop signals. Given the 64 possible codon combinations, this genetic code exhibits redundancy, allowing several synonymous codons to encode for the same amino acid. Historically, synonymous codons were perceived as being inconsequential to the resultant protein structure due to the consistent amino acid they encode for, termed a ‘silent’ mutation [17]. However, recent revelations in biochemistry and bioinformatics challenge this viewpoint.

The phenomenon of optimal codons, the specific codon most frequently used by an organism, showcases the rich variability across evolutionary lineages [17, 18]. The variance in codon usage bias alone enables accurate prediction of an organism’s phylogenetic lineage and the organelle of origin for a genetic sample [19]. This intricate codon usage can be attributed to metabolic demands, regulatory mechanisms governing gene expression, and adaptive responses within the organism. This complexity is further demonstrated by the fact that synonymous codons employ different tRNAs. Distinct tRNAs are recruited for different codons, meaning that organisms favoring a select set of optimal codons can economize on their tRNA production to optimize metabolic efficiency and potentially offer adaptive advantages in specific ecological niches [20, 21, 22, 23, 24].

Moreover, studies have elucidated that silent mutations can indeed influence the final protein structure [18, 25]. The same amino acid sequence, when encoded by different synonymous codons, can produce structurally diverse protein domains [25]. One postulated mechanism attributes this variance to the rate of translation; rare tRNAs can slow down or even pause translation, providing the emerging polypeptide chain an extended window to attain its native conformation, possibly even leveraging the ribosome as a chaperone [18, 26, 27, 28]. Conversely, abundant tRNAs expedite translation. Either scenario could be potentially optimal for a protein, given its specific requirements. Additionally, optimal codons are thought to translate with higher accuracy compared to their lower usage counterparts [22]. Therefore, we hypothesize that the codon sequence of a protein likely contains more information than the amino acid sequence.

If codon sequences are truly superior from an information standpoint, then it follows that pLMs based on codon sequences should provide an increased depth of protein knowledge. To test if a pLM based on codon sequences can discern additional information, we developed cdsBERT (CoDing Sequence Bidirectional Encoder Representation Transformer). cdsBERT was seeded with ProtBERT and further trained on 4 million CoDing Sequences (CDS) compiled from the NIH and Ensembl databases. The resulting model was subsequently trained via a modified student-teacher Knowledge Distillation (KD); we termed this vocabulary extension pipeline MELD (Masked Extended Language Distillation). Our hypothesis was that a shift in synonymous codon embeddings within the TEM would indicate a nontrivial addition of protein information after applying MELD. Furthermore, we also validated our model by conducting Enzyme Commission (EC) classification with cdsBERT vs. ProtBERT to establish how intuitively the codon latent space maps to an example of protein function.

## 2 Methods

### 2.1 Coding sequence compilation

The CDS is a DNA sequence that determines the sequence of amino acids in a protein [29]. We utilize two curated sources of CDS in our research. First was the NIH Consensus CDS (CCDS) project, which contains a high-quality aggregate of CDS over the entire mouse and human genomes [29]. To introduce wider phylogenetic diversity into our dataset, we compiled the CDS data of over 300 additional genomes from Ensembl [30] as a second source. Preprocessing included removing sequences that 1) did not start with ATG or end in a traditional stop codon, 2) were longer than 1000 amino acids (computational efficiency), and 3) had lengths that were not multiples of three. To optimize data storage, we used a novel codon single letter code shown in **Table 1**, to reduce dataset storage space by one-third. We attempted to make the letter code as relevant as possible, keeping the human optimal codons as their corresponding uppercase amino acid code, and the second most frequent synonymous codons in the lowercase version. Other seemingly random additions were kept as close to the original amino acid letter as possible. We mapped methionine to “(” and the stop codons to either “)”, “}”, or “]”. This made all possible open reading frames easily visible by containing them within different forms of brackets.

**Table 1:**
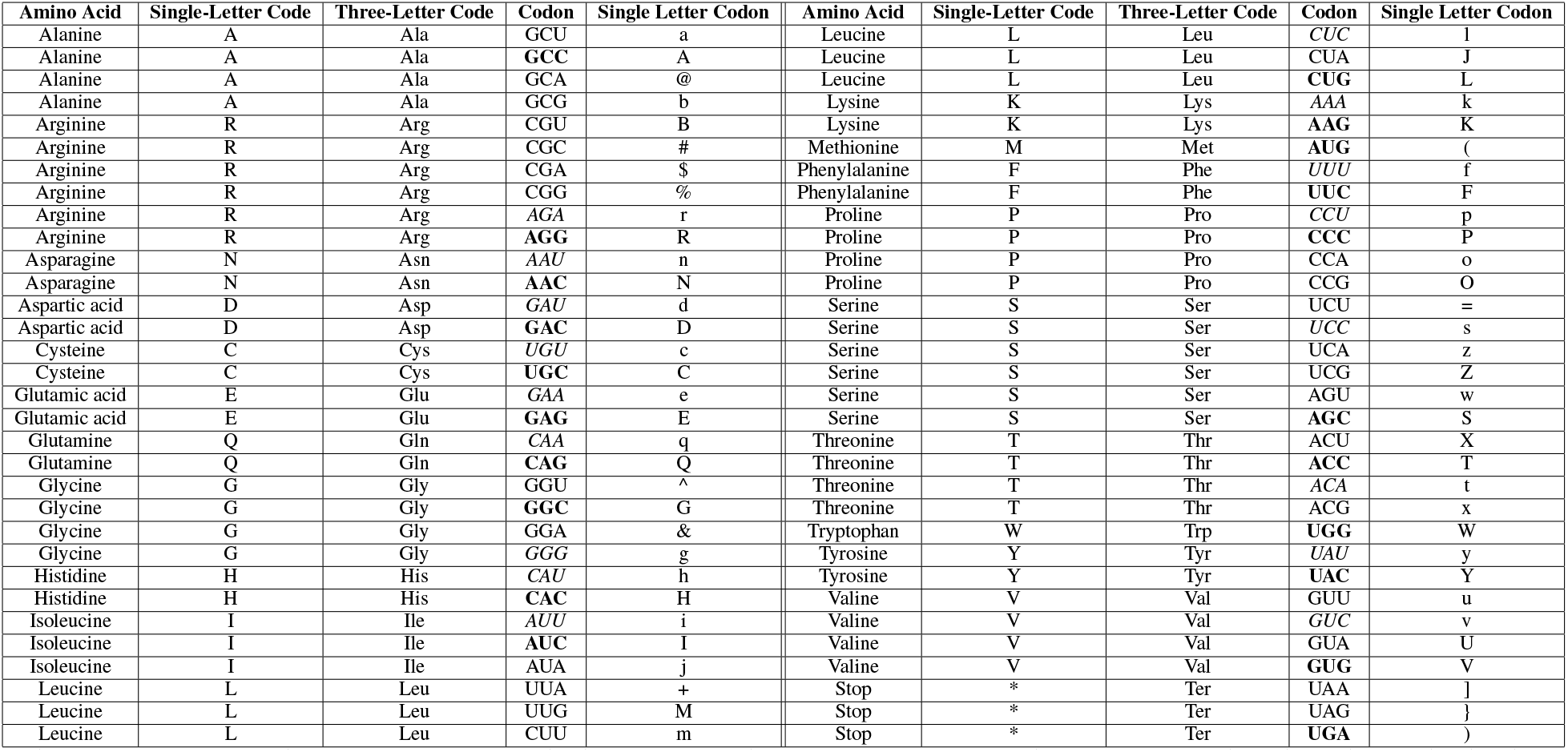
Translation between amino acid identifiers and novel single letter identifiers. Bold codons have the highest frequency in humans and the italicized codons have the second highest.

### 2.2 Protein language models

pLMs, underpinned by transformer neural network architectures, have demonstrated great utility in protein science. Originally designed for natural language processing (NLP) tasks, transformers have remarkable proficiency in interpreting a multitude of sequential data; including protein, nucleotide, and broader chemical sequences [1, 12].

A cornerstone of this proficiency lies in the token embeddings, a representation method that transforms words in natural language, or amino acids in protein sequences, into numerical vectors. In essence, each amino acid is analogous to a word in natural language and is mapped to a unique integer, while the entire protein sequence can be viewed as a sentence. These integers then serve as indices, providing access to a predefined matrix or lookup table, the TEM. As such, the TEM is a learned vector representation that encapsulates the syntactic and semantic nuances of each amino acid [1, 31].

Central to the transformer architecture is the multi-head self-attention mechanism. This mechanism is adept at capturing long-range dependencies between tokens in a sequence. It achieves this by evaluating the importance of each token relative to every other token in the sequence. The mathematical representation of this self-attention can be formulated as:

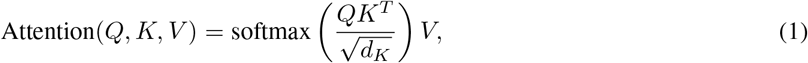

where *Q, K*, and *V* denote the query, key, and value matrices, respectively. *d*_*k*_ is the dimension of the key matrix. These matrices are extracted from the embeddings of amino acids, described by:

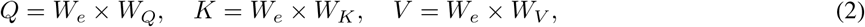

with *W*_*e*_ being the TEM and *W*_*Q*_, *W*_*K*_, and *W*_*V*_ representing learned weights [31].

Further expanding on the self-attention mechanism, the multi-head attention subdivides the input sequence, creating multiple sets of query, key, and value matrices. Each of these sets, or “heads”, processes the sequence independently, capturing unique relationships and patterns. These individual outputs are then consolidated to provide a comprehensive understanding of the sequence with a further learned linear projection [31].

In total, a single “transformer layer” is made up of a combination of attention and feed-forward layers, which introduce non-linear transformations in the latent space. Typically, many of these transformer layers need to be stacked together to effectively capture semantic information in the embeddings and contextual information with attention. In our case, we utilized a pre-trained model: ProtBERT-BFD.

ProtBERT-BFD is a pLM with 30 transformer layers, 16 attention heads, an embedding dimension of 1024, and trained on the Big Fantastic Dataset (BFD) comprised of over 2 billion amino acid sequences in a self-supervised fashion [12]. We chose to use this specific pLM due to its modest size at 420 million parameters compared to much larger counterparts. The size and excellent protein understanding enabled efficient model use and training. Additionally, the 1024 embedding size is quite large relative to its total parameter count. This means that the capability of this model in capturing semantic and nuanced synonymous codon information is intuitively higher than a model with a smaller embedding dimension in which to store that information.

### 2.3 MELD

To effectively extend the vocabulary of a pLM we developed a novel pipeline using TEM seeding, masked language modeling (MLM), and student-teacher knowledge distillation (KD) called MELD.

Firstly, instead of training a TEM from scratch for cdsBERT we expanded the TEM of ProtBERT from size 30 (normal amino acids, special amino acids, special tokens) to 69, to include the 64 codons and 5 special tokens [32]. Instead of initializing weights randomly we seeded all synonymous codons with the same weights from their corresponding amino acid. Stop codon token embeddings, which do not have a corresponding amino acid, were seeded randomly.

After TEM seeding we utilize MLM to learn new vocabulary specific semantic information. MLM is the process of randomly hiding portions of the text and then having the model “fill in the blanks,” repeating this many times [33]. The MLM task was formatted as the following objective with a corpus of tokens *U* = *u*_1_, …, *u*_*n*_ maximizing the likelihood:

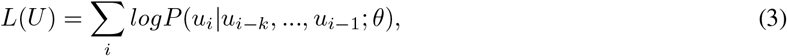

where *k* is the context window of masked tokens [33], and the conditional probability *P* is predicted by cdsBERT with parameters *θ* (*W*_*e*_, *W*_*Q*_, *W*_*K*_, *W*_*V*_, etc. ∈ *θ*). The output distribution of tokens was learned through the hidden or latent space outputs *H* = *h*_0_, …, *h*_*m*_:

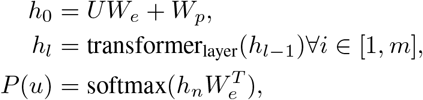

where *U* = (*u*_−*k*_, …, *u*_−1_) is the context vector of tokens, *m* is the number of transformer layers, and *W*_*p*_ is the position embedding matrix [33]. *L*(*U*) was maximized during training by minimizing the cross-entropy between the model output and the true label.

To learn more fine-grain and contextual information we utilized KD after MLM; a methodology that uses a contrastive loss to compare the latent output of two models with the goal of improving the representation of the smaller or less trained model [34, 35]. Codon sequences were fed into cdsBERT with the equivalent amino acid sequences into a state-of-the-art pLM while aligning their numerical interpretations in space. The pLM we chose was the base Ankh model; a highly efficient pLM that offers a state-of-the-art yet open-source biochemically-relevant latent space representation despite its modest size [13]. We kept Ankh frozen and in half-precision with cdsBERT in half-precision as well for expedited training.

To build an effective vector representation for contrastive loss we averaged the last hidden state over the length for BERT and Ankh variants. Additionally, we utilized a trainable linear layer to project the 768-dimensional vector from Ankh to a 1024-dimensional vector for the loss function. This way, we could ensure the cdsBERT latent space somewhat copied the biochemical relevance of Ankh with the freedom to move weights in a larger vocabulary.

We chose a custom contrastive loss to enable the use of information from within the batch to reward the model when a sample’s corresponding pair was closest in space and move embeddings away from samples that were not matching pairs [36]. The combination of batch-wise matrix multiplication and softmax had the form:

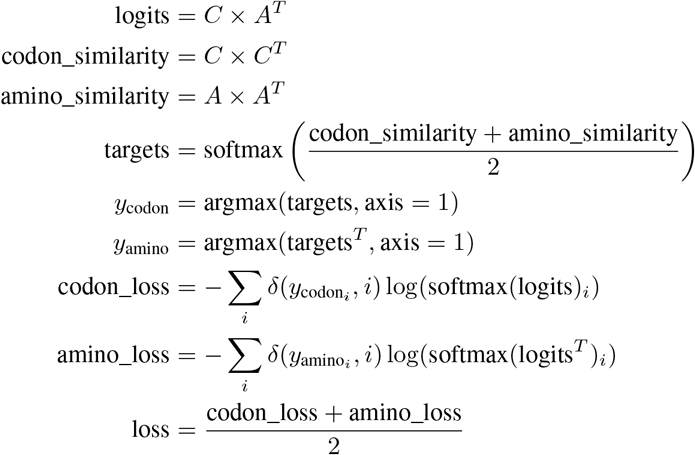

where *C* was the batched codon representation, *A* was the amino acid representation, conducted one-hot encoding, and *n* was the batch size. Using this loss strategy, we contrasted the BERT models after MLM with Ankh base using 500,000 total sequences from a random subset of our CDS data. The final models were denoted with a +; cdsBERT+ and ProtBERT+ for comparison.

### 2.4 Enzyme commission number prediction

EC numbers serve as a comprehensive and hierarchical numbering scheme to classify and describe enzyme-catalyzed reactions based on their generalized function [37, 38]. This structured classification begins at a broad level and progressively narrows to specific enzyme actions. Primary sequence data has fueled successful EC predictions using sophisticated deep-learning techniques including pLMs [39, 40, 41, 42, 43, 44]. Thus, we used EC prediction as a gold-standard to discern biochemically-relevant protein representations.

For EC data acquisition, we used human and mouse samples from the CCDS database. Preliminary analysis showed that CCDS typically matched exactly to their UniProt counterpart while Ensembl sequences tended to be slightly different. To overcome this, CCDS were matched to their UniProt counterparts, then trimmed by sequences with a single EC number annotation whose length did not exceed 1000 amino acids. The CCDS codon sequences were processed using cdsBERT, while their amino acid counterparts were subjected to ProtBERT for comparative analysis.

To prepare feature vectors for classification, we input the corresponding sequences to their respective model and averaged across the length. After the full dataset had been encoded, each sample *v* was normalized via the mean of the set *μ* and standard deviation 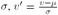, resulting in a *v*^*′*^ used for classification. In addition to *v*^*′*^, we incorporated two prominent classification algorithms believed to be particularly efficacious for data that is hypothesized to be well-separated: Support Vector Machines (SVMs) and K-Nearest Neighbors (KNN). SVMs operate by finding a hyperplane that best divides a dataset into classes, making it suitable for high-dimensional and particularly separable data [19, 45]. KNN is a non-parametric method that classifies a data point based on how its neighbors are classified, proving effective for datasets where class boundaries are irregular [45]. We include KNN for comparison as SVM has a small chance of resulting in a particularly good or bad fit. This shortcoming was further alleviated by cross-validation (CV) where independent fits were used to optimize hyperparameters for the classifiers.

Labels were associated with their respective vectors either as the first EC digit or as the full class mapped to a unique integer. We looked at first and full EC digits to give a reference frame to our resultant metrics. First EC digit is a much easier less discriminative task and thus would result in higher metrics. Thus this comparison lets us conclude if the end performance was data-limited given the preprocessed EC CCDS dataset size of 2629. Because of the small dataset size, we reported the average 10-fold CV metrics as our evaluation metrics. The training and evaluation process used is illustrated in **Figure 1**.

**Figure 1:**
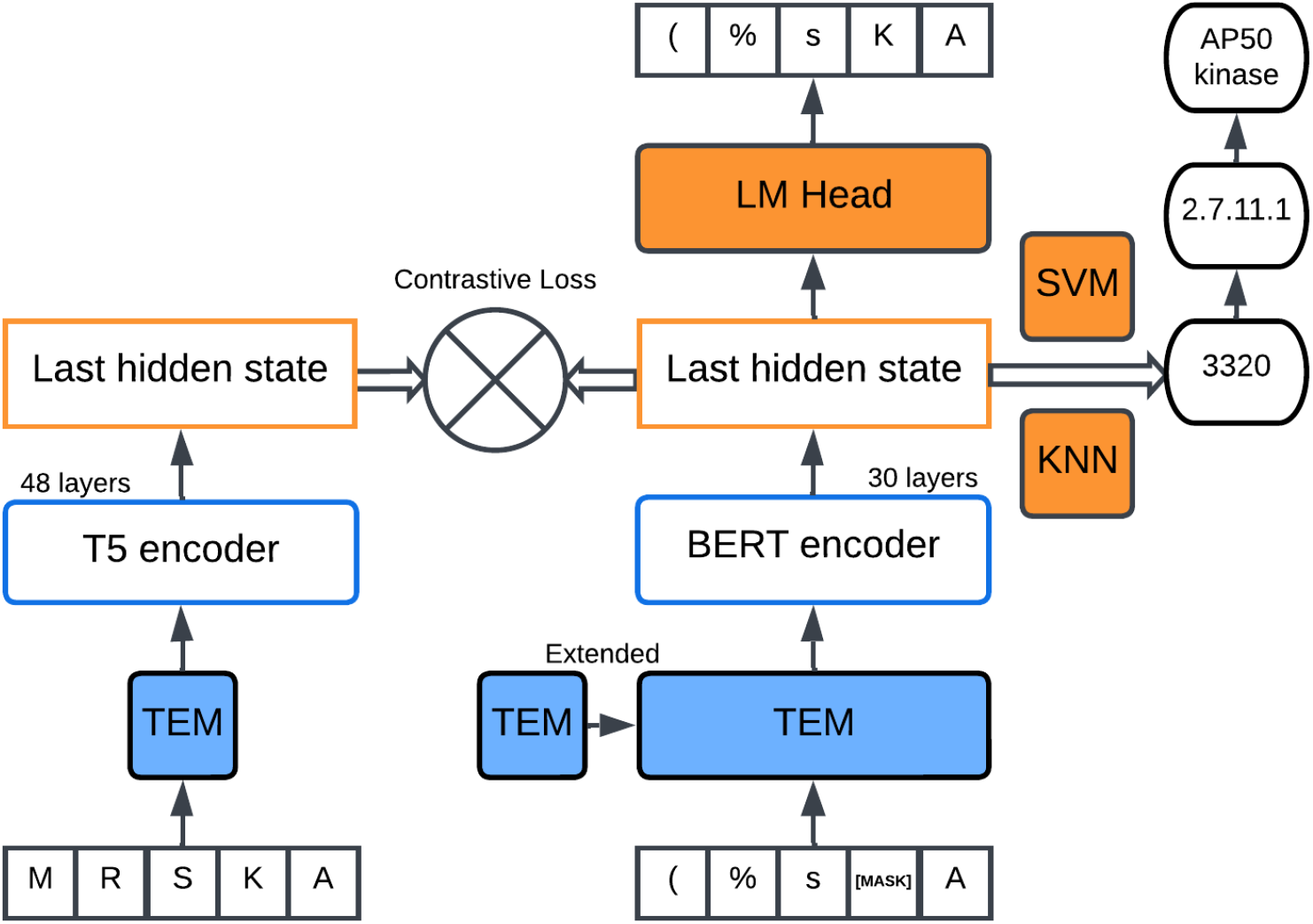
Model diagram used for the MELD process and model evaluation. Stage 1: seed cdsBERT with ProtBERTBFD and extend the TEM with the codon vocabulary. Synonymous codons started with their corresponding amino acid ProtBERT embedding, stop codons were randomly initialized. Stage 1 used MLM on codon sequences from CCDS and Ensembl. Stage 2: KD with Ankh-base. Vector embeddings from the averaged last hidden state were used to train KNN and SVM classifiers for EC number prediction.

### 2.5 cdsBERT weight analysis

To characterize changes to the synonymous codon embeddings after training, we used a Principal Component Analysis (PCA) plot to visualize the approximate magnitude of any movement in the reduced projected space of two principal components [46].

To compare the movement of the codon embeddings, we used a novel vector comparison of the form:

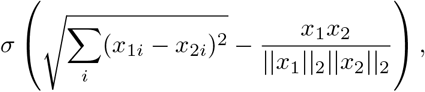

where *σ* is the sigmoid function, *x*_1_ and *x*_2_ are token embeddings from two different codons, and *║x*_1_ *║*_2_ and *║x*_2_*║*_2_ are the *L*2 norms of *x*_1_ and *x*_2_, respectively. By subtracting the cosine similarity from the Euclidean distance, we quantified the angle and magnitude difference of the vectors. The sigmoid function normalized the measure between 0 and 1 for easy comparison, enabling a standardized vector difference metric that incorporates magnitude and angle in space. We refer to this metric as the sigmoid-distance-similarity. We calculated the sigmoid-distance-similarity between every codon and their start location in the TEM. Stop codons were excluded from this analysis as they were initialized randomly. We refer to this as the *movement list*.

To determine embedding movement correlations we compared the movement list to three lists derived from the codon usage dataset [19]:

- *Average usage list*: The average codon usage across diverse phylogeny, preserving generalized conserved codon usage mechanisms.
- *Kingdom ranking list*: A list of codons and a corresponding score depicting how influential that codon’s frequency is in predicting the phylogenetic identity of a sample.
- *DNA ranking list*: A list of codons and a corresponding score depicting how influential that codon’s frequency is in predicting the DNA type (origin) identity of a sample.

Because the feature ranking analysis of the CUF dataset constructed two individual rankings based on lasso and random forests, we needed a unified heuristic to construct a single list. We extended the feature ranking by using a strategic voting scheme to combine feature ranking outputs from the lasso, random forests, extreme gradient boosting, mutual information, and recursive feature elimination. The result of the ensemble was a single list of codons and scores for kingdom ranking and DNA ranking.

We aimed to decipher potential relationships between the movement list and three distinct variables: average usage, kingdom, and DNA lists. To robustly determine the nature and strength of these associations, we employed both Pearson’s and Spearman’s correlation analyses.

During our training stages, we were curious how MLM and KD affected the weights more broadly than just the token embeddings during vocabulary extension. To analyze this, we measured the average MSE between these weight types in ProtBERT vs. cdsBERT to look at MLM and cdsBERT vs. cdsBERT+ to look at KD.

## 3 Results

In the initial stages, cdsBERT’s performance, following MLM, lagged behind the base ProtBERT model; cdsBERT achieved an accuracy of 62.4% versus ProtBERT’s 74.7% on full EC numbers using KNN, as detailed in **Table 2**. Given the substantial computational demands of MLM, and with our dataset’s extensive size (approximately 9.5 million sequences) it was not feasible to complete even a single epoch of training. This led us to surmise that cdsBERT might be undertrained with its new vocabulary, resulting in an inferior latent space representation.

**Table 2:**
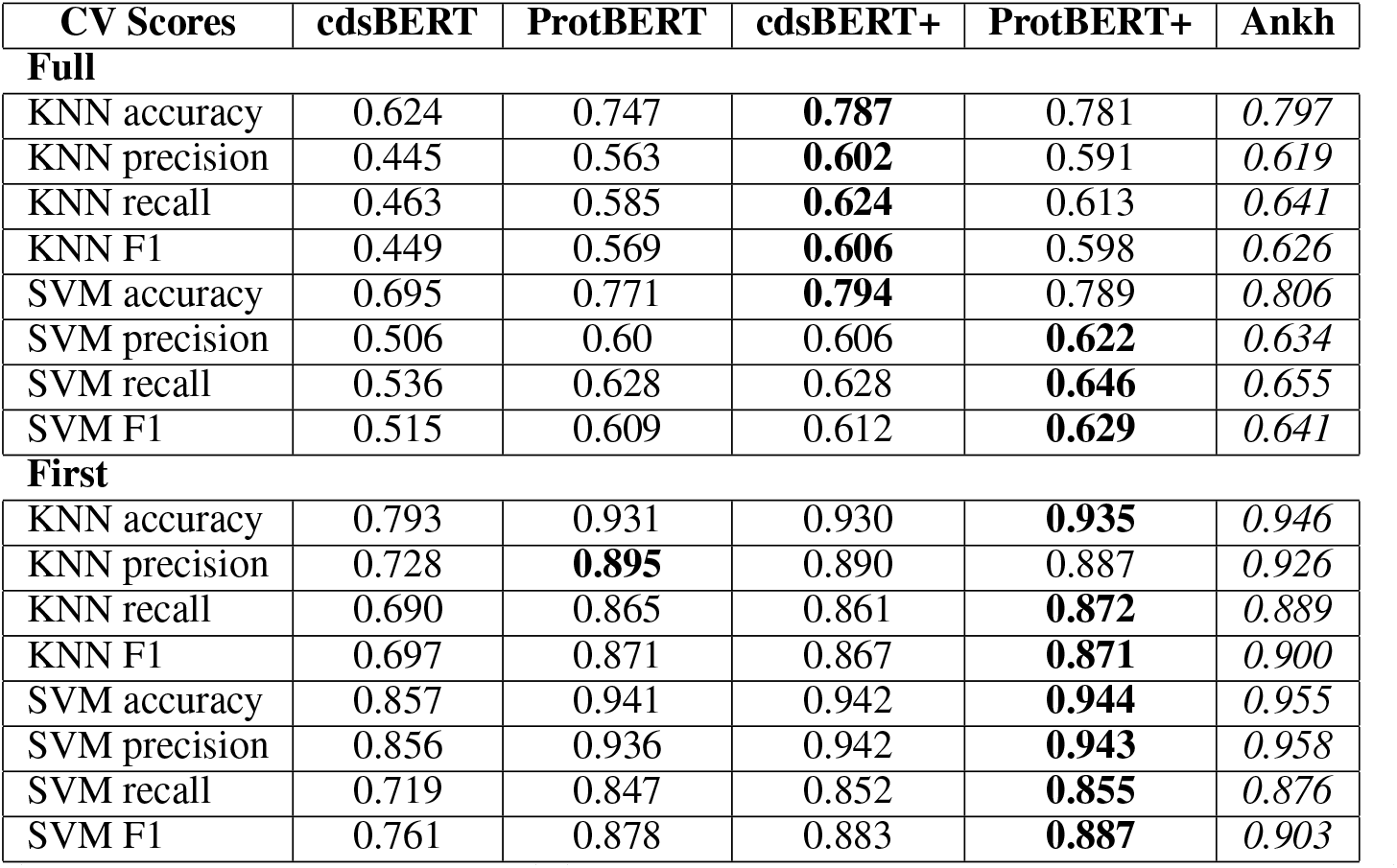
EC prediction of using KNN, SVM and the protein vector representations from various models. Reported metrics are averaged from 10-fold cross validation. Results for full EC numbers mapped to unique integers are on top while the results for just predicting the first EC digit are on the bottom. The best performing result from our work is bolded. The initial cdsBERT after MLM performed poorly after MLM but increased dramatically after KD training, surpassing the original ProtBERT weights. Most importantly, cdsBERT+ performed slightly higher than ProtBERT+ on full EC numbers. Results for Ankh-base are shown for comparison, italicized because Ankh performs better than our architecture variants.

| CV Scores | cdsBERT | ProtBERT | cdsBERT+ | ProtBERT+ | Ankh |
| --- | --- | --- | --- | --- | --- |
| **Full** |  |  |  |  |  |
| KNN accuracy | 0.624 | 0.747 | **0.787** | 0.781 | *0.797* |
| KNN precision | 0.445 | 0.563 | **0.602** | 0.591 | *0.619* |
| KNN recall | 0.463 | 0.585 | **0.624** | 0.613 | *0.641* |
| KNN F1 | 0.449 | 0.569 | **0.606** | 0.598 | *0.626* |
| SVM accuracy | 0.695 | 0.771 | **0.794** | 0.789 | *0.806* |
| SVM precision | 0.506 | 0.60 | 0.606 | **0.622** | *0.634* |
| SVM recall | 0.536 | 0.628 | 0.628 | **0.646** | *0.655* |
| SVM F1 | 0.515 | 0.609 | 0.612 | **0.629** | *0.641* |
| **First** |  |  |  |  |  |
| KNN accuracy | 0.793 | 0.931 | 0.930 | **0.935** | *0.946* |
| KNN precision | 0.728 | **0.895** | 0.890 | 0.887 | *0.926* |
| KNN recall | 0.690 | 0.865 | 0.861 | **0.872** | *0.889* |
| KNN F1 | 0.697 | 0.871 | 0.867 | **0.871** | *0.900* |
| SVM accuracy | 0.857 | 0.941 | 0.942 | **0.944** | *0.955* |
| SVM precision | 0.856 | 0.936 | 0.942 | **0.943** | *0.958* |
| SVM recall | 0.719 | 0.847 | 0.852 | **0.855** | *0.876* |
| SVM F1 | 0.761 | 0.878 | 0.883 | **0.887** | *0.903* |

To ascertain whether the suboptimal cdsBERT performance was due to inadequate training or inherent to the new vocabulary, we applied KD to cdsBERT, producing cdsBERT+, and similarly applied KD to ProtBERT, yielding ProtBERT+. The results were promising: cdsBERT+ accuracy increased by over 16.3% to 78.7%, outperforming the original ProtBERT. In contrast, ProtBERT+ improved only 3.4% to 78.1%. Notably, cdsBERT+ beat ProtBERT+ in all KNN metrics and SVM accuracy. Further KD training with either Ankh-base or Ankh-large did not enhance cdsBERT+ or ProtBERT+ performance. This implies that both models may have approached their performance ceilings on our EC dataset, potentially constrained by their architecture and size.

For the first EC digits, cdsBERT+ and ProtBERT presented comparable results, with cdsBERT+ excelling in SVM performance and ProtBERT in KNN. However, ProtBERT+ slightly outperformed both. The full results are in **Table 2** with Ankh-base added for comparison. We used the average last hidden state instead of the [CLS] for an aggregate representation in training and evaluation as a fair comparison because when using the [CLS] token or pooler output with ProtBERT it performed significantly worse. The pooler output used was standard from the Hugging Face transformers package, a dense linear layer and the Tanh activation function [32]. We are unsure exactly why ProtBERT is not robust to this addition, but random or previously learned initialization of the pooler output severely degraded ProtBERT’s performance on EC prediction after KD. The pooler output with cdsBERT performed nearly the same as the averaging scheme.

Our model weight MSE analysis (**Figure 2**) offers insight into cdsBERT’s significant performance leap upon transitioning to cdsBERT+ (**Figure 2B**). We hypothesized that the MLM phase would predominantly refine the TEM responsible for semantic information, whereas KD would optimize attention layers for contextual learning. Our results confirmed this to an extent. During the vocabulary extension in MLM, the TEM underwent more extensive updates relative to other layers, based on the training we conducted (**Figure 2A**). In contrast, KD primarily influenced the linear layers, followed by the TEM and attention layers (**Figure 2B**). These findings underline the overall efficacy of MLM and KD in enriching semantic and contextual understanding during vocabulary expansion in pLM vocabulary using MELD. Whereas these have been used together in previous language models for the creation of distilled models, we observed significantly greater effects of KD following MLM.

**Figure 2:**
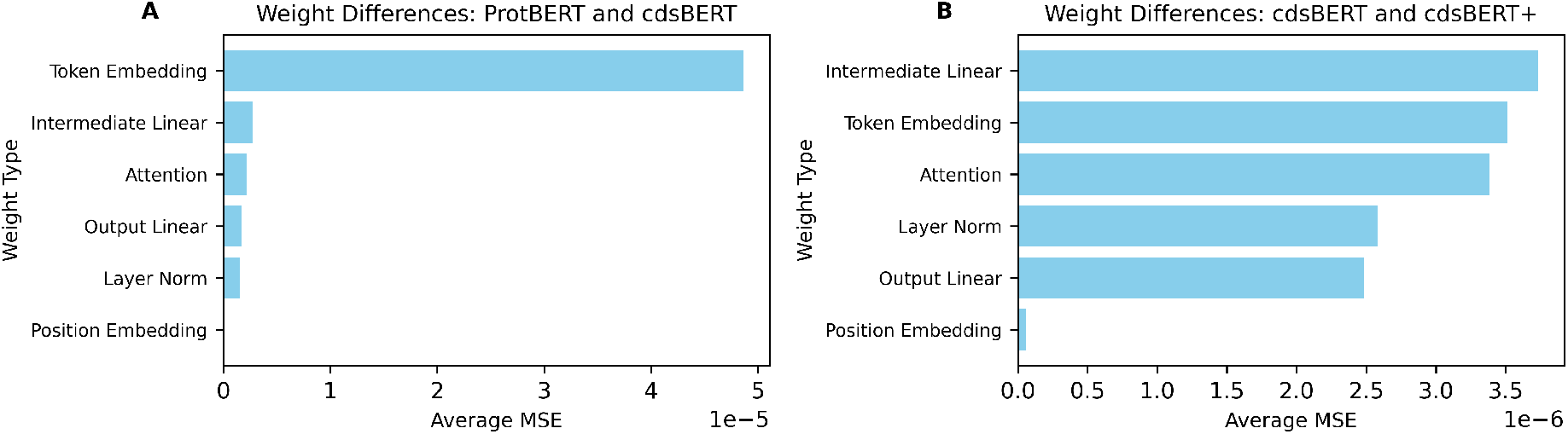
**(A)** MSE measurements of cdsBERT seeded with ProtBERT after MLM illustrates that the main update during MLM was of the TEM, learning semantic information. **(B)** The same analysis between the MLM and KD weights (cdsBERT+) shows that while the TEM was updated during KD, much of the fine-tuning was broadly in the space of contextual understanding via intermediate linear layers and attention.

A deeper dive into the weight modifications revealed intriguing dynamics. Upon training, synonymous codons exhibited distinct shifts, clustering around their native amino acid embeddings from ProtBERT. The PCA plot (**Figure 3AB**) presents this in a 2D space, where codons are labeled based on their corresponding amino acid identity and physiochemical attributes. Positive amino acid groupings are clustered tightly along PC1 while negative ones are clustered along PC2. Neutral and polar amino acids are mostly placed on the lower half of PC2 with polar amino acids are largely constrained to the right side of PC1. Importantly, the three stop codons have clustered together despite being initialized randomly. Although synonymous codon shifts appear slight within the full 2D PCA projection, closer examination of the six synonymous serine codons (**Figure 3A** inset), for example, reveals unique groupings and deviations among the codons, suggesting unique semantic information and possible usage patterns have emerged from the CDS samples.

**Figure 3:**
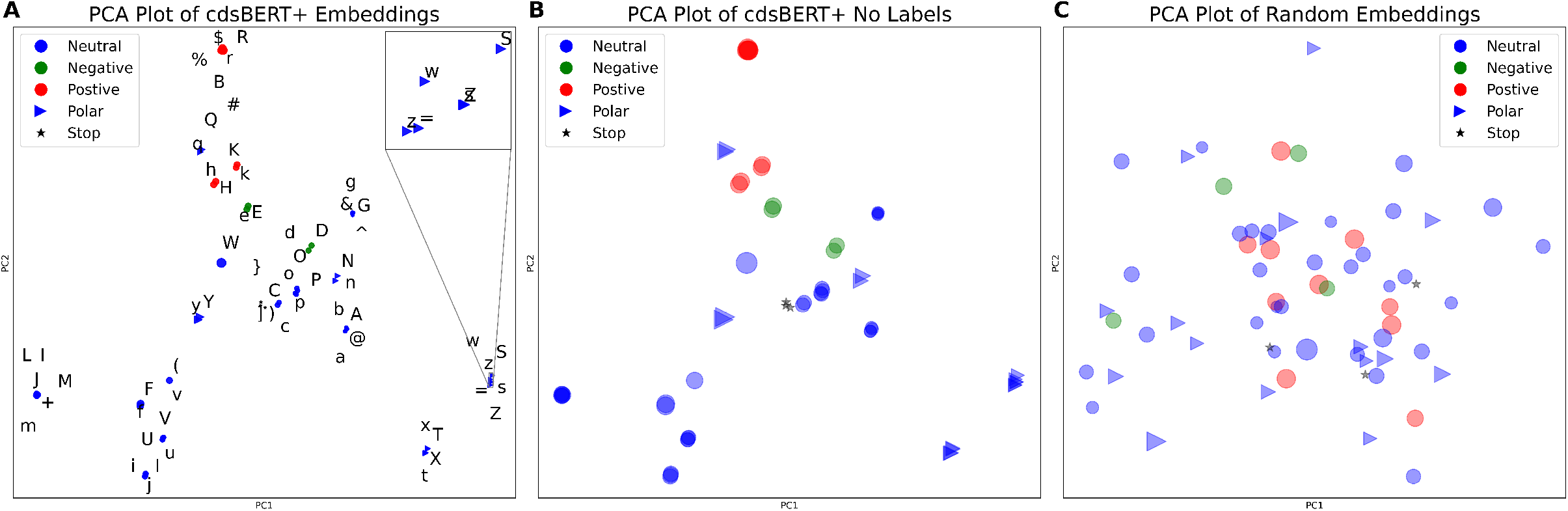
**(A)** PCA plot illustrating the token embeddings of cdsBERT+ projecting from 1024 to two-dimensional space. Codons are grouped according to their corresponding amino acid properties. **(B)** Same as **A** but without label clutter, larger points, and added opacity for easy visualization. **(C)** Randomized embeddings are shown to get a sense of how well the groupings are. Codon embedding points in all subplots are scaled by the weight of their corresponding amino acids in Daltons.

To comprehensively quantify codon shifts within the token embedding space, we calculated the sigmoid-distance-similarity for every pair of embeddings (**Figure 4**). In this distogram, purple represents a larger shift distance while orange denotes a closer distance. Each element on the plot corresponds to the metric applied between the *i*th and *j*th codon embedding. The diagonal, showing a perfect similarity, indicates that a codon embedding is perfectly similar to itself. Along the diagonal, 21 distinct regions are evident, with each corresponding to synonymous codons. Notably, this representation strongly suggests that synonymous codons evolved independently during the training phase; if they had moved in unison each region would be a consistent dark orange shade.

**Figure 4:**
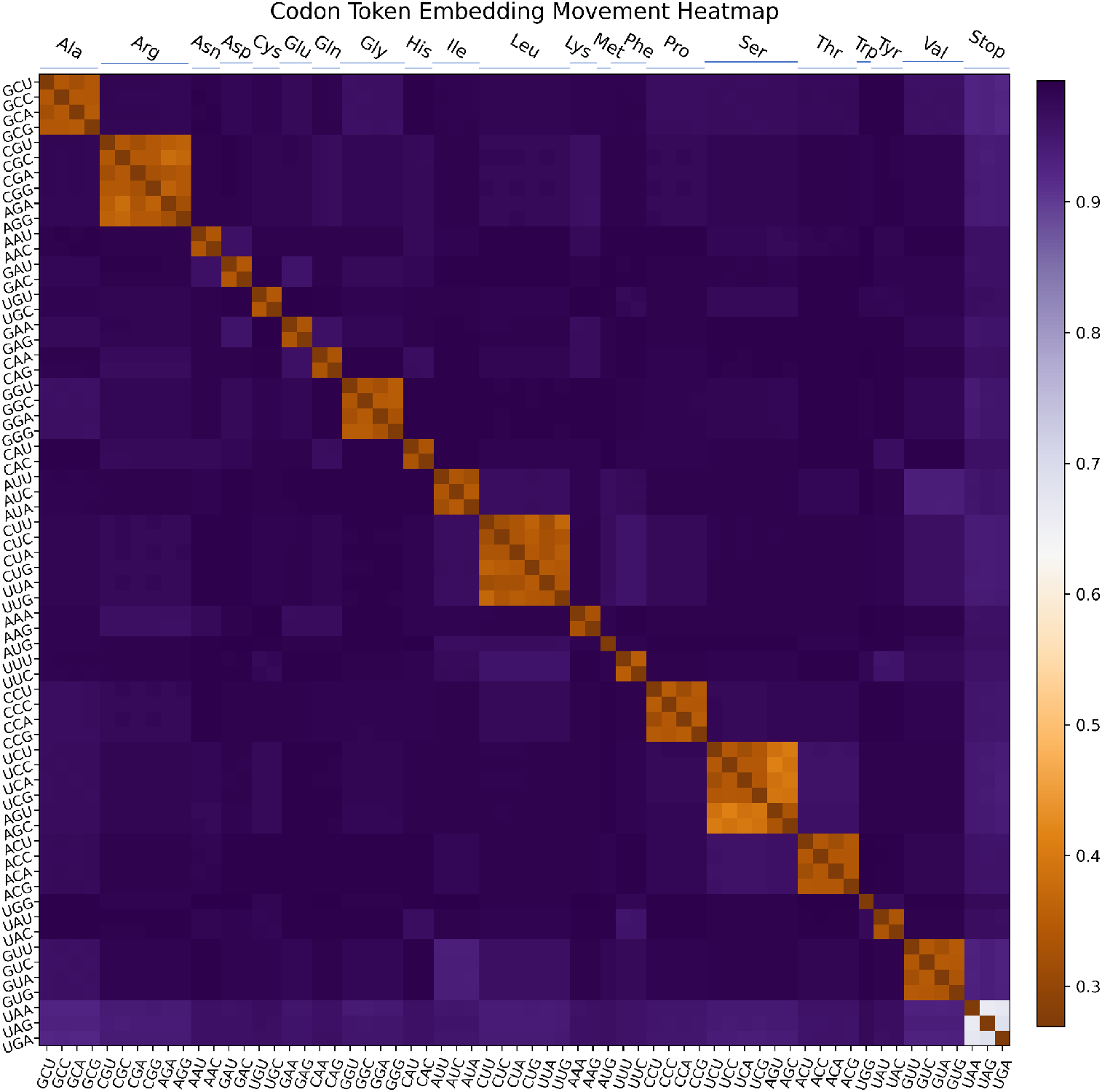
Distogram comparing the distance of each codon embedding to every other one, using the sigmoid-distancesimilarity metric. Purple signifies greater distance and orange is closer. Each entry is the metric applied between the ith and jth codon embedding, where the diagonal is perfect similarity between one codon embedding and itself. Around the diagonal are 21 distinct segments, each corresponding to synonymous codons.

To interpret potential causes that underlie the movement of synonymous codons, we turned to a previously published codon usage frequency (CUF) dataset [19]. This dataset contains each of the 64 codon frequencies of a nucleotide sample (summing to 1) and corresponding labels of species, relative phylogenetic identity, and organelle of origin. With approximately 13,000 CUFs across wide phylogeny, this dataset enabled correlation analysis of our codon embeddings to potentially explain movement patterns throughout MELD. We used the kingdom-set of the CUF data to split the phylogenetic classes into virus, bacteria, archaea, animals, and plants, and DNA type classes into nuclear, mitochondrial, and chloroplast [19]. Trends of the codon shifts in movement, kingdom ranking, DNA ranking, and average usage, as defined by Pearson’s or Spearman’s correlations are shown in **Figure 5**. Notably, a medium negative correlation existed between codon movement and its average usage, suggesting codons used more frequently were less prone to shifting away from the native amino acid embedding during training. This relationship is intuitive, as the higher frequency codons should most resemble their native amino acid counterparts since amino acids in sequences are preferentially from this subset.

**Figure 5:**
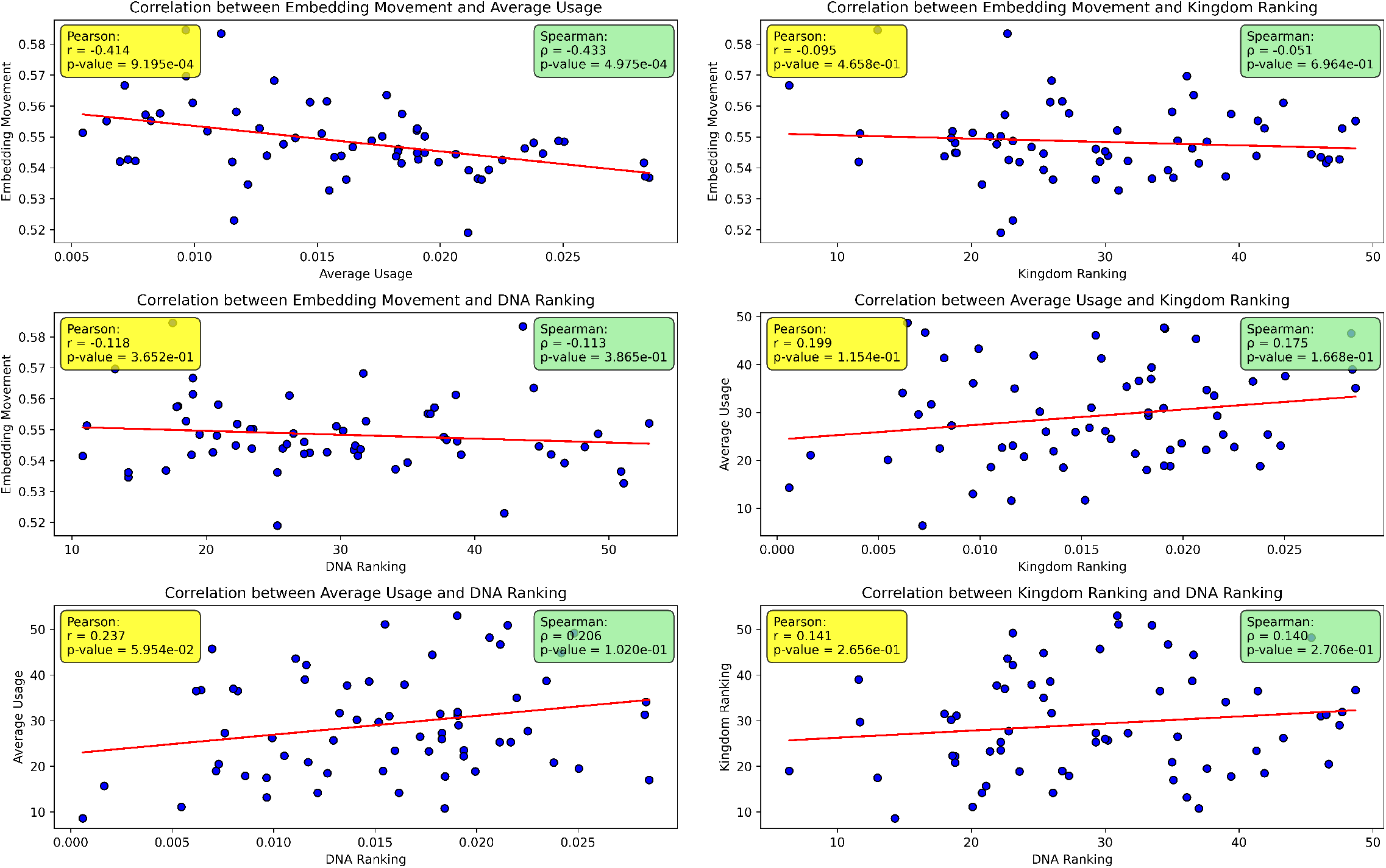
Statistical correlation measurements between all combinations of reported codon phenomena. The four lists of codons and corresponding measures are embedding movement, average usage over phylogeny, feature ranking for phylogenetic prediction, and feature ranking for DNA type prediction.

The only other statistically significant correlation was the small positive relationship between average usage and DNA ranking. Thus, the more likely a codon is to be used the more likely it is differentially used in different organelles, such as the nucleus, mitochondria, and chloroplast. These two statistically significant correlations imply that there should be a relationship between the embedding movement and codon influence on DNA type prediction, but this relationship is weak and not significant. The other weak correlations are also intuitive, namely 1) usage and influence on kingdom prediction were positively correlated, 2) influence on DNA and kingdom were positively correlated, and 3) embedding movement and kingdom ranking were effectively not correlated.

## 4 Discussion

Recent studies into codon usage bias have cast light on its significance, especially in scenarios where the fundamental structure of the resulting protein is impacted. This work accentuates the necessity for codon awareness in achieving pLMs that are truly biologically relevant. cdsBERT is emblematic of this exploratory step in pLM vocabulary, steering toward a more informed and enriched language model.

To establish cdsBERT, we demonstrated a novel and efficient pipleine for extending the vocabulary of pLMs termed MELD. We suspect that by leveraging new semantic and contextual information from MLM and KD natural language models could be compared and extended for greater proficiency in multiple languages even without intuitive TEM seeding. Furthermore, the potential for other chemical language model extensions where there is translatable nomenclature is evident.

For full EC number prediction, the addition of the codon vocabulary increased the overall performance, even when ProtBERT was further trained with KD. However, as these performance gains were small, we only expect a slight performance increase from switching vocabularies. Despite codon usage alone affecting post-transcriptional modifications, mRNA structure, protein structure, influencing the speed of translation, and enabling phylogenetic and DNA type prediction, the vast majority of protein information can be summarized with the amino acid vocabulary. Of course, in the limit, a well-trained pLM would not be hindered by the codon vocabulary as the model could learn to ignore the codons and simply translate to amino acids if information encoded within codons was truly never helpful. A well-trained model could even ignore or not ignore this additional information depending on the context needed. Therefore, whereas the precise amount of information added by codon sequences over amino acid counterparts is yet unknown, we conclude there is an addition of information.

The correlation analysis used to understand possible reasons for codon movement in the embedding space yielded intuitive but mostly negligible relationships. However, the correlation between average codon usage and codon embedding movement was highly statistically significant with a medium negative relationship (**Figure 5**). This showcases the ability of cdsBERT to capture extremely generalized codon-based information due to its exposure to vast phylogeny. Meaning, the general trends in usage, when averaged across phylogeny, are at a minimum captured in the TEM. This finding is exciting, demonstrating potential for a well-trained codon-based pLM holding general trends within the TEM and contextualizing sequence properties based on the codon usage of a given input. This is similar to previous applications of codon bias based on single nucleic acid samples instead of the averages of whole genomes, for example, horizontal gene transfer and open reading frame prediction [19]. With this in mind, we believe the correlations with kingdom and DNA ranking would increase dramatically if cdsBERT was fine-tuned on a specific phylogeny, learning more specific usage patterns instead of generalized ones. Thus, this correlation analysis has potential in serving as a metric to determine if a codon-based model has been fine-tuned effectively.

Our studies highlight the pressing need for meticulous CDS mappings to expansive protein sequence repositories, such as UniProt. This would enable the creation of vast sequence databanks conducive to self-supervised learning, as well as curated datasets for evaluating the biochemical significance of the derived latent spaces. Looking forward, the momentum generated by this research underscores the importance of forging a foundational model based on extended vocabularies. Such models, drawing inspiration from successful endeavors like ESM or Ankh, could set the benchmark for future pLMs.

Given an adequate data pool, a codon-centric foundation model could undergo fine-tuning akin to contemporary pLMs. This could encompass a spectrum of annotation tasks, ranging from EC and gene ontology categorization, to predicting structures and interactions. On the side of generative models, possibilities span areas like protein design. Moreover, a foundation model’s versatility could be stretched to consider organism-specific codon biases, particularly in industrial contexts for protein and peptide synthesis; enabling more efficient and effective industrial translation [47, 48].

In conclusion, cdsBERT, reiterates the profound potential of utilizing codon awareness. By emphasizing the balance between fundamental amino acid information and the nuanced codon details, we hope to build awareness for the addition and use of CDS data in large protein repositories.

## 5 Data and model availability

The data used for training is available through public databases ncbi.nlm.nih.gov/projects/CCDS/ and ensembl. org/info/data/biomart. The weights of cdsBERT+ are available through Hugging Face at huggingface.co/ GleghornLab/cdsBERT.

## 6 Funding and acknowledgments

This work was partly supported by the University of Delaware Graduate College through the Unidel Distinguished Graduate Scholar Award (L.H.), the Ronald E. McNair Post Baccalaureate Achievement Program (N.R.), and the National Institute of Allergy and Infectious Diseases R21AI157889. Any opinions, findings, and conclusions or recommendations expressed in this material are those of the authors. We would like to acknowledge the valuable feedback and critical review of this work from Katherine Nelson Ph.D. and Krithika Umesh. Figures were generated with Lucidchart and Matplotlib.

